# Cryo-EM structures of mycobacterial MCC reveal carrier-domain translocation between catalytic sites

**DOI:** 10.64898/2026.03.23.713199

**Authors:** Ajit Yadav, Bogdan I. Florea, Sebastian Geibel

**Author notes:** **Materials & correspondence** Correspondence and requests for materials should be addressed to Sebastian Geibel.

## Abstract

3-Methylcrotonyl-CoA carboxylase (MCC) is a biotin-dependent enzyme complex that catalyzes an essential step in leucine degradation. In mycobacteria, MCC is formed by AccA1 and AccD1, but its structural organization has remained poorly defined. Here, we report two high-resolution cryo-EM structures of the endogenous, biotin-bound α_6_β_6_ complex from *Mycobacterium smegmatis*. The MCC architecture comprises a central hexameric carboxyltransferase (β/CT) core flanked by trimeric biotin carboxylase (α/BC) modules. The structures capture BC- and CT-engaged conformations, in which the biotin carboxyl carrier protein (BCCP) engages either the BC or CT active site, providing structural evidence for long-range carrier-domain translocation. Conformational rearrangements within the BC module suggest a role in regulating BCCP engagement. Together, our findings provide insights into carrier-domain dynamics in mycobacterial MCC.

## Introduction

Branched-chain amino acid degradation contributes to central carbon metabolism and metabolic flexibility in bacteria [1]. In mycobacteria, leucine catabolism proceeds through 3-methylcrotonyl-CoA carboxylase (MCC), a biotin-dependent enzyme complex that catalyzes the ATP-dependent carboxylation of 3-methylcrotonyl-CoA to 3-methylglutaconyl-CoA [1-3]. Biotin-dependent acyl-CoA carboxylases share a modular architecture in which catalysis proceeds through two sequential steps at spatially separated active sites. In the first step, the biotin moiety attached to the biotin carboxyl carrier protein (BCCP) is carboxylated by the biotin carboxylase (BC) domain in an ATP- and Mg^2+^-dependent reaction [4,5], using bicarbonate as the carboxyl donor. The carboxylated biotin is then shuttled by the flexible BCCP domain to a distinct carboxyltransferase (CT) active site, where the carboxyl group is transferred to the substrate [6,7].

In mycobacteria, MCC is formed by the AccA1-AccD1 complex [8]. AccA1 comprises the BC and BCCP domains, with an additional BC-CT interaction (BT) domain, whereas AccD1 provides the CT active site. Catalysis requires long-range shuttling of the biotinylated BCCP domain between spatially separated BC and CT active sites, coupling ATP-dependent carboxylation to substrate modification.

High-resolution structures of MCC holocomplexes from bacteria and eukaryotes have established a conserved α_6_β_6_ dodecameric architecture with a characteristic α_3_β_6_α_3_ organization, in which a central carboxyltransferase core is flanked by biotin carboxylase-BCCP modules [9-13]. However, structural information remains limited to a small number of bacterial (*Pseudomonas*) and eukaryotic systems (*Homo sapiens, Trypanosoma, Leishmania*), and MCC complexes from mycobacteria have been largely unexplored. Structural evidence for mycobacterial MCC has been restricted to low-resolution electron microscopy data, which confirmed the overall architecture but lacked the resolution to define domain organization, active-site architecture, and catalytically relevant conformations [8]. During the preparation of this work, a preprint reported a cryo-EM structure of the MCC holo-complex from *Mycobacterium smegmatis*, in which the BCCP domain is positioned at the CT region [14]. Here, we report two high-resolution cryo-EM structures of the AccA1-AccD1 MCC complex, defining its molecular architecture and capturing distinct conformations with BCCP engaged at either the BC or CT active site.

## Results

### Purification and identification of the endogenous AccA1-AccD1 holoenzyme

During StrepTactin affinity purification, we unexpectedly co-purified the endogenous MCC complex consisting of the biotin carboxylase subunit AccA1 and the carboxyltransferase subunit AccD1 from cytosolic and membrane fractions of *Mycobacterium smegmatis* (Supplementary Table 1). The AccA1-AccD1 complex was further purified to homogeneity by size-exclusion chromatography (Fig. 1a).

**Fig.1.**
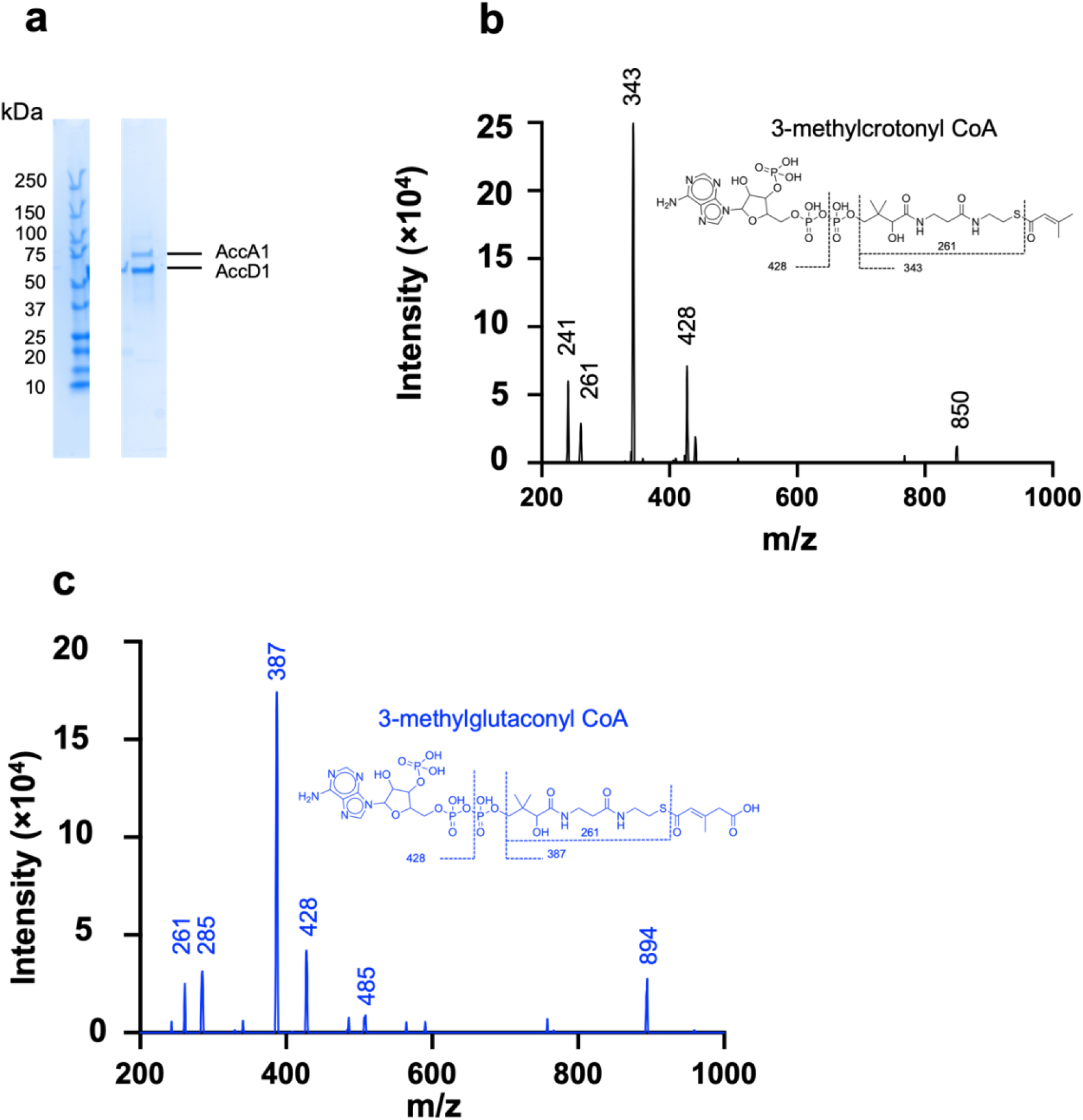
Purification and Biochemical Characterization of MCC complex. **a**, SDS-PAGE analysis of the endogenous 3-methylcrotonyl-CoA carboxylase purified from *Mycobacterium smegmatis*. **b**, MS/MS spectrum (precursor ion scan of m/z=850, in positive ion mode) and chemical structure of 3-methylcrotonyl-CoA. **c**, MS/MS spectrum (precursor ion scan of m/z=894, in positive ion mode) and chemical structure of 3-methylglutaconyl-CoA.

### Purified MCC is catalytically active

To assess catalytic activity, we employed LC-MS/MS to directly monitor product formation. In positive ion mode, precursor ions were detected at m/z 850 for 3-methylcrotonyl-CoA and m/z 894 for the carboxylated product, 3-methylglutaconyl-CoA (Fig. 1b,c). Both species exhibited a characteristic fragment at m/z 428 corresponding to the adenosine diphosphate moiety. Additional fragments corresponding to the acyl-CoA moiety were observed at m/z 343 for 3-methylcrotonyl-CoA and m/z 387 for 3-methylglutaconyl-CoA, consistent with efficient carboxylation of the substrate. Together, these data demonstrate that the purified MCC complex is catalytically active toward 3-methylcrotonyl-CoA.

### Architecture of the mycobacterial MCC complex

To define the molecular architecture of mycobacterial MCC, we analyzed the purified AccA1-AccD1 complex from both membrane and cytosolic fractions by single-particle cryo-electron microscopy. Ab initio reconstruction, followed by heterogeneous and non-uniform refinement [15], yielded maps at overall resolutions of 2.2 Å and 2.9 Å, respectively (Supplementary Fig. 1-2, Supplementary Table 2), both revealing a canonical MCC assembly with D3 symmetry. The BC module, in the CT-engaged BCCP state, could only be modeled partially.

The AccA1-AccD1 holoenzyme assembles into a 748 kDa heterododecamer comprising six α-subunits (AccA1) and six β-subunits (AccD1), organized in a four-layered α_3_β_6_α_3_ architecture. Six AccD1 protomers form a hollow, barrel-shaped hexamer composed of two stacked trimeric rings, constituting the central carboxyltransferase (CT) core. On each face of the β_6_ ring, three AccA1 subunits assemble into trimeric layers that form the biotin carboxylase (BC) modules, thereby completing the α_6_β_6_ holoenzyme (Fig. 2a,b).

**Figure 2.**
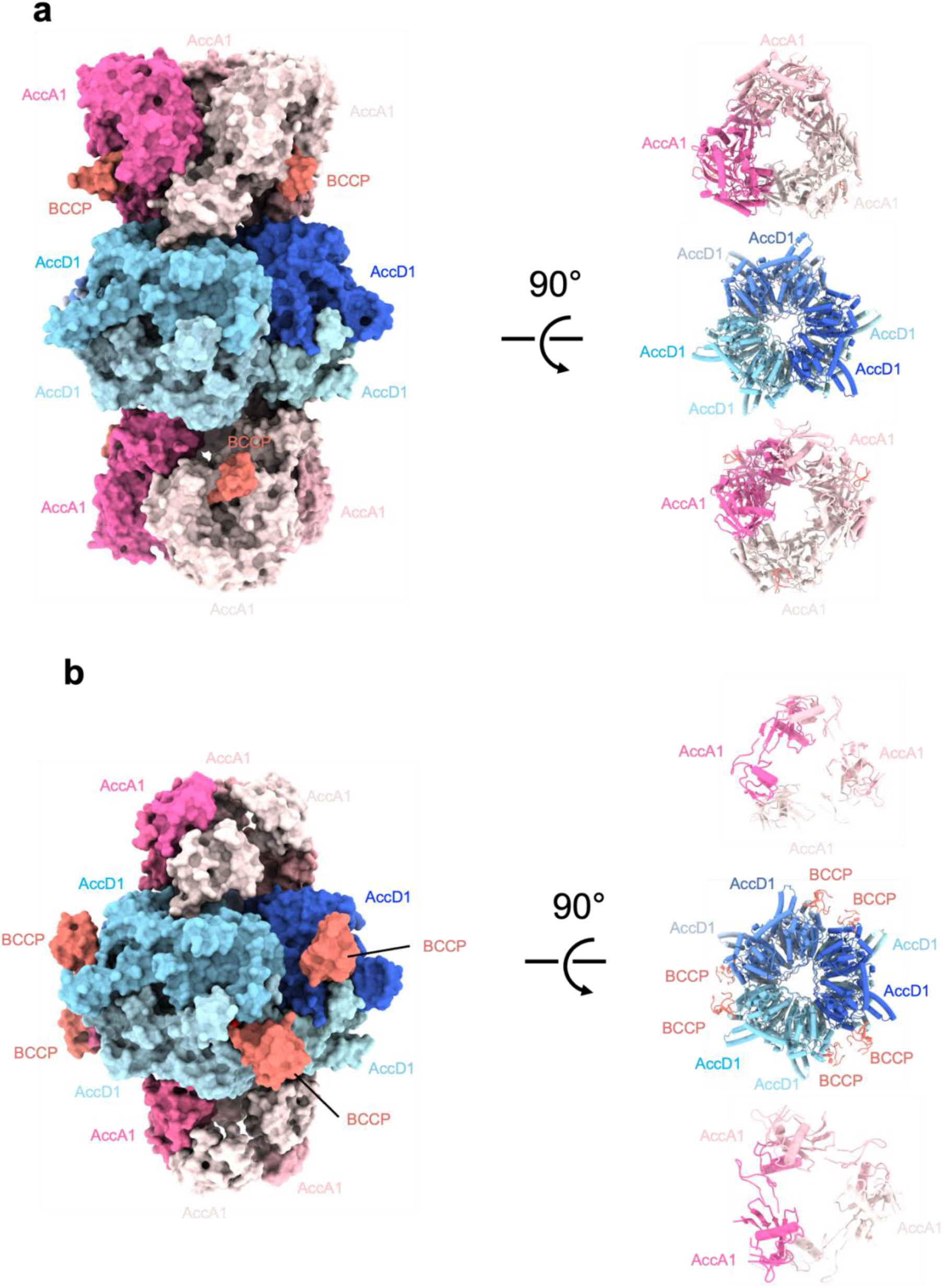
Architecture of the MCC complex in BC-engaged BCCP and CT-engaged BCCP states. Atomic models of the MCC complex (surface representation) built into composite cryo-EM maps (Supplementary Fig. 1,2) showing the BC-engaged. **a**, and CT-engaged **b**, BCCP states. The central carboxyltransferase (CT/β) core is flanked by two biotin carboxylase (BC/α) modules positioned at the top and bottom. BCCP domains (salmon) engage either the BC modules (BC-engaged state) or relocate toward the CT core (CT-engaged state). A 90° rotated view (cartoon representation) shows the CT/β subunits (AccD1) forming a hexameric core (shades of blue) organized as a dimer of β trimers, while the BC/α subunits (AccA1) assemble as trimers (shades of pink).

The BT domain mediates the primary interface between AccA1 and the AccD1 CT core, anchoring the BC-BCCP module to the central hexamer. An extended loop within the BT domain engages the CT subunit through hydrogen-bond interactions involving conserved residues W486, G491, W492, and R493 with residues G53, S54, F56, and L493 of AccD1. In addition, R493 establishes a salt bridge with D47 of AccD1 (Fig. 3a). This interaction positions the flexible BCCP domain for long-range shuttling between the BC and CT active sites.

**Fig. 3.**
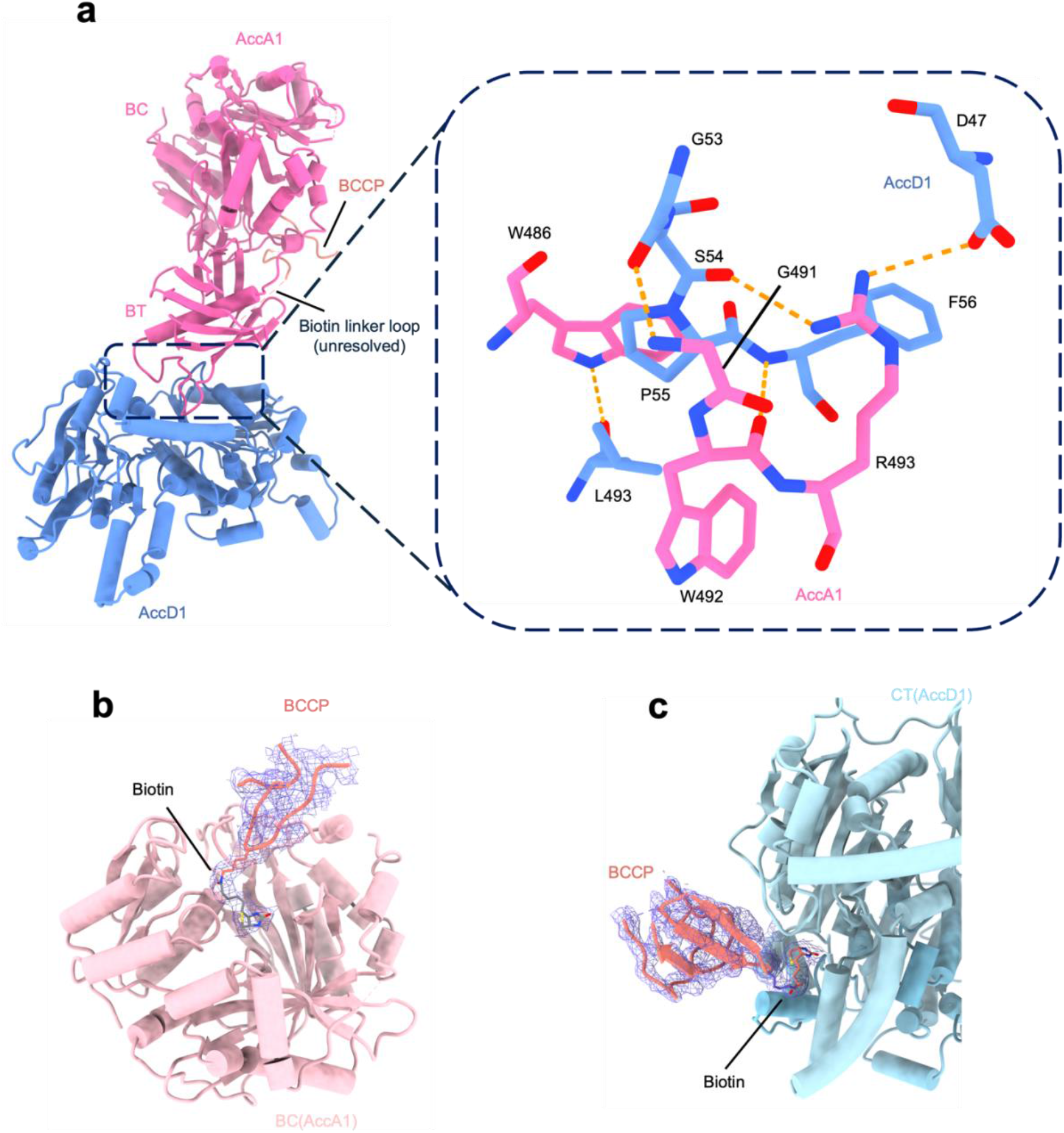
Structural details of α-β subunit interactions and distinct active-site engagement of BCCP. **a**, Left, cartoon representation of the interaction region between the α and β subunits. The extended loop (residues 482-499) in the BT domain interacts with all domains in the β subunit. Right, detailed view of the interacting residues shown in stick representation. Yellow dashed lines indicate hydrogen bonds and salt bridge interactions. **b**, BC-engaged BCCP (blue mesh density), with biotinylated Lys628 (blue mesh density) positioned within the ATP-binding pocket of AccA1 (Pink). **c**, CT-engaged BCCP (blue mesh density) transiently samples the substrate-entry region of AccD1 (light blue).

The overall organization of the AccA1-AccD1 holoenzyme closely matches MCC architectures reported in other organisms, confirming the conserved quaternary arrangement of biotin-dependent acyl-CoA carboxylases (Supplementary Fig. 3). Consistent with this, structural superposition of the carboxyltransferase subunit AccD1 with previously reported AccD1 structures from *M. tuberculosis* and the human carboxyltransferase subunit MccB reveals two conserved alanine-glycine-based oxyanion holes that are required for carboxyl transfer (Supplementary Fig. 4).

### BCCP adopts distinct catalytic positions at BC and CT active sites

In the two cryo-EM structures, the BCCP domain occupies distinct positions corresponding to different catalytic states. In one structure, the biotinylated BCCP domain is located at the active site of the BC module, with the uncarboxylated biotin oriented toward the nucleotide-binding pocket, consistent with a carboxylation-competent state (Fig. 3b). In the second structure, the BCCP domain is positioned at the CT active site, placing the uncoraboxylated biotin in proximity to the substrate-binding pocket, consistent with a CT-engaged conformation that may represent a post-carboxyltransfer or substrate-sampling state (Fig. 3c). The two states are separated by a large-scale displacement of the BCCP domain of approximately 61 Å between the BC and CT active sites (Fig. 4a). This movement is enabled by the flexible linker connecting the BCCP domain to the BT domain, allowing the carrier domain to shuttle between spatially separated catalytic centers. Together, these structures illustrate long-range carrier-domain translocation within MCC required for catalytic activity.

**Fig. 4.**
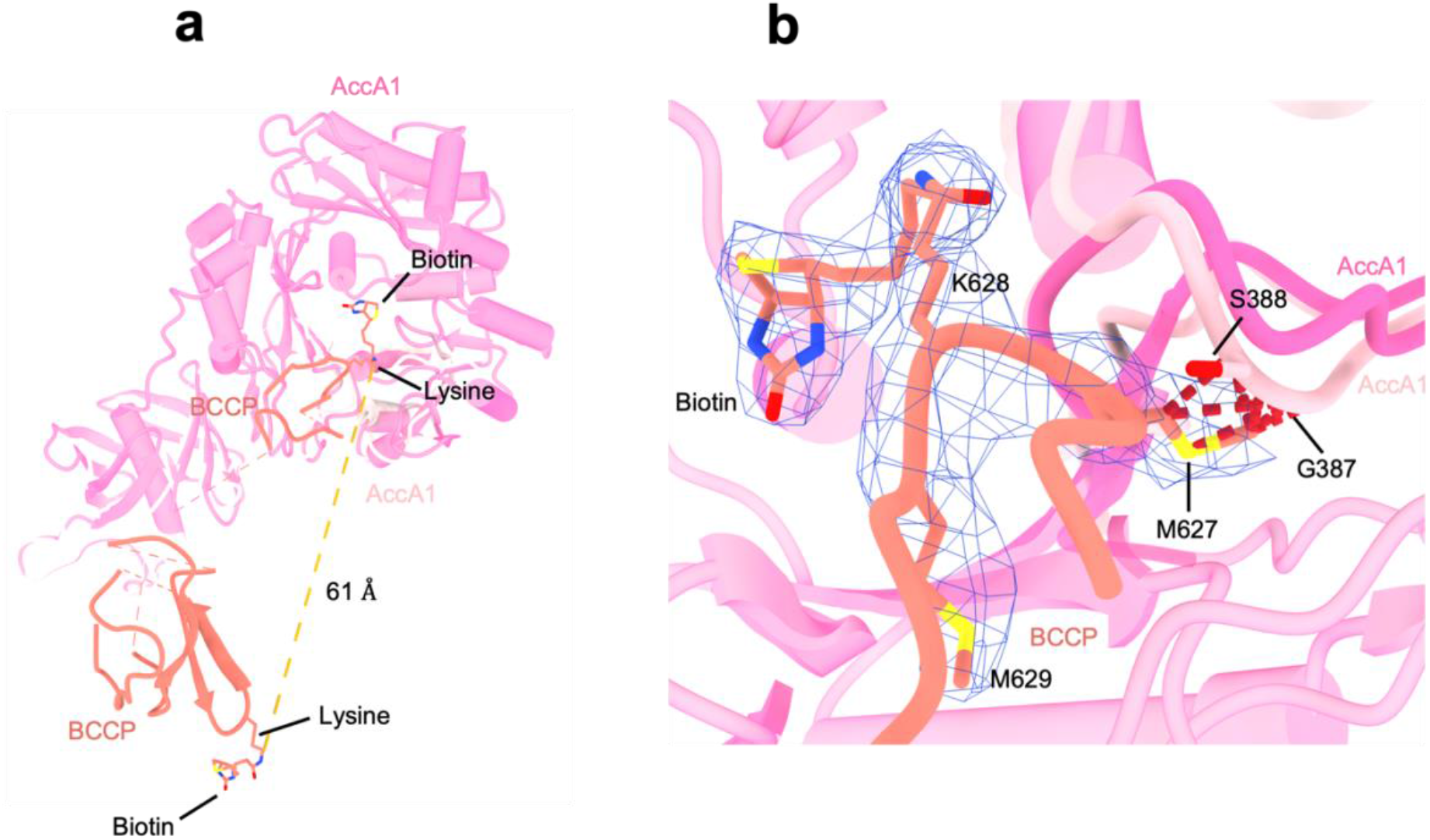
Conformational changes associated with BCCP movement. **a**, Distinct positions and conformations of BCCP (salmon) in the BC-engaged and CT-engaged states of the MCC complex. The CT subunit is omitted for clarity. Biotinylated lysine are shown in stick representation. **b**, Structural superimposition of the BC domain in the two states: BC-engaged BCCP (dark pink) and CT-engaged BCCP (light pink). The red dot indicates steric clashes between Met627 of the conserved biotin-binding motif (blue EM density) and residues Gly387 and Ser388 of the BC domain in the CT-engaged BCCP state of MCC complex.

### BCCP-mediated conformational changes in the MCC complex

Comparison of the BC- and CT-engaged states reveals a conserved loop in the BC domain (residues 384-392) that undergoes a conformational rearrangement (Supplementary Fig. 5). In the BC-engaged state, this loop accommodates the biotin-binding motif (M627-K628-M629) of the BCCP domain. In contrast, in the CT-engaged state, residues G387 and S388 shift toward the biotin-binding site, resulting in a steric clash with residue M627 of BCCP (Fig. 4b). This rearrangement is therefore incompatible with BCCP engagement at the BC active site. In the BC-engaged state, this loop accommodates the biotin-binding motif (M627-K628-M629) of the BCCP domain. In contrast, in the CT-engaged state, residues G387 and S388 shift toward the biotin-binding site, resulting in a steric clash with the conserved residue M627 of BCCP (Fig. 4b). This rearrangement is therefore incompatible with BCCP engagement at the BC active site.

In contrast to human MCC, where loop rearrangements in the CT module have been proposed to preclude BCCP binding [13], no significant conformational differences are observed within the CT subunit between the two states (Supplementary Fig. 6).

## Discussion

In this study, we define the molecular architecture of the endogenous 3-methylcrotonyl-CoA carboxylase (MCC) from *M. smegmatis*. Our cryo-EM structures reveal a canonical α_6_β_6_ dodecameric assembly with a four-layered α_3_β_6_α_3_ organization, consistent with MCC architectures reported in other systems [9-13], but previously unresolved at high resolution in mycobacteria [8]. A central finding of this work is the visualization of two distinct conformations in which the BCCP domain engages either the BC or the CT active site. These states are separated by a large displacement of the carrier domain, providing structural evidence for long-range domain translocation during MCC catalysis.

Our data further identify a conserved loop in the BC domain that undergoes a conformational rearrangement between the two states. In the CT-engaged conformation, this loop adopts a position that is incompatible with BCCP binding at the BC active site, suggesting a role in coordinating access of the carrier domain. This mechanism contrasts with human MCC, where rearrangements in a loop within the CT module have been proposed to regulate BCCP engagement [13]. The absence of comparable conformational changes in the CT subunit of mycobacterial MCC suggests that coordination of carrier-domain positioning may be mediated primarily at the BC domain rather than the CT module.

Together, these observations suggest that the CT core in mycobacterial MCC functions as a structurally stable scaffold that accommodates carrier-domain movement without requiring major conformational remodeling. The differences observed between mycobacterial and human MCC indicate that distinct strategies have evolved to coordinate carrier-domain engagement in different systems.

## Material and Methods

### Bacterial strains and growth conditions

*Mycobacterium smegmatis* mc^2^155 was transformed with the plasmid *pMyNT* empty vector and pMyNT: Esx3_strep_PPE4, in which the Esx3_strep_PPE4 construct is expressed under the control of the acetamidase promoter. Cultures were grown in Luria-Miller (LB) broth supplemented with 0.2% (v/v) glycerol at 180 rpm. Induction was performed with 0.2% (w/v) acetamide at an optical density at 600 nm (OD_600nm_) of 0.5 and cells were grown until a final optical density of 2.0-2.1. Cells were pelleted by centrifugation and washed in 3× PBS buffer. For the control, uninduced cultures of pMyNT empty vector were processed in parallel using identical conditions.

### Purification of MCC complex from membrane and soluble fraction

Cell pellet from all three different conditions (induced *pMyNT::Esx3_strep_PPE4, pMyNT* empty vector, and uninduced *pMyNT* empty vector) were resuspended at a 1:5 (w/v) ratio in buffer A (300 mM NaCl, 50 mM HEPES, 1 mM DTT, pH 8.0) supplemented with EDTA-free protease inhibitors (Roche) and lysozyme (Roche). Cells were lysed by three passages through an Emulsiflex-C3 homogenizer (Avestin), and unlysed debris was removed by centrifugation at 10,000 × g for 15 min at 4 °C. The membrane fraction was isolated by ultracentrifugation (45,000 rpm, 1 h, 45Ti rotor, 4 °C). The soluble fraction was loaded directly onto a pre-equilibrated Strep-Tactin™ XT 4Flow™ column (IBA), washed with buffer A until a stable UV baseline was reached, and eluted with buffer B (buffer A supplemented with 50 mM biotin, pH 8.0). Membranes were solubilized in buffer A containing 10× CMC OG (Anatrace) overnight at 4 °C. Insoluble material was removed by ultracentrifugation (45,000 rpm, 1 h, 45Ti rotor, 4 °C), and the solubilized membrane fraction was applied to a pre-equilibrated Strep-Tactin™ XT 4Flow™ column. Columns were washed with buffer A containing 5× CMC DM and eluted with buffer B supplemented with 5× CMC DM. Soluble and membrane eluates were subjected to size-exclusion chromatography on a Superose 6 Increase 10/300 GL column (GE Healthcare) equilibrated in buffer C (300 mM NaCl, 20 mM HEPES, 1 mM DTT, pH 8.0), with 2× CMC DM added for membrane samples. Eluted fractions were analyzed by SDS-PAGE followed by Colloidal Blue staining.

### Liquid chromatography-mass spectrometry (LC-MS)

Chromatographic analysis of 3-methylcrotonyl-CoA was carried out using ion-pairing reverse-phase high-performance liquid chromatography (IP-RP-HPLC) on an Agilent 1260 Infinity II system (Agilent Corporation, Santa Clara, CA, USA) with a NUCLEODUR EC C18 column (50 × 4.6 mm, 3 μm particle size). The chromatographic separation was performed at a flow rate of 1 mL min^−1^ using a gradient elution program. Initially, the column was equilibrated with mobile phase A consisting of 90% H_2_O and 10% (v/v) 100 mM ammonium acetate. A 10 μL sample was injected and maintained under these starting conditions for 1.5 min. The solvent composition was then gradually changed over 9 min to mobile phase B composed of 90% acetonitrile (CH_3_CN) and 10% (v/v) 100 mM ammonium acetate. After reaching mobile phase B, the column was rinsed with this solvent for 2 min. Subsequently, the system was returned to mobile phase A through a 10 s gradient, followed by an additional 2 min equilibration period before the next sample injection. Mass spectrometric detection was performed on an Agilent 6475 LC/TQ system (Agilent Technologies, Santa Clara, CA, USA) equipped with an AJS electrospray ionization (ESI) source, operated in positive ion mode. The ion source parameters were set as follows: nebulizer pressure 20 psi, drying gas flow 10 L min^−1^, drying gas temperature 100 °C, and capillary voltage 4000 V. Data acquisition was carried out within a mass range of m/z 200-1000. Fragmentor voltage and collision energy values were optimized to obtain maximal signal intensity and efficient fragmentation of 3-methylcrotonyl-CoA.

### *In vitro* MCC activity assay

MCC enzymatic activity was confirmed by detection of carboxylated 3-methylcrotonyl-CoA using ion-pairing reverse-phase high-performance liquid chromatography (HPLC) coupled with triple quadrupole mass spectrometry (TQ-MS/MS). Enzymatic reactions were performed in a 25 μL reaction mixture containing 50 mM HEPES buffer (pH 7.2), 2 mM ATP, 8 mM MgCl_2_, 50 mM NaHCO_3_, and 50 μM 3-MCC substrate. Reactions were initiated by adding the purified MCC complex to a final concentration of 0.50 μM, followed by incubation at 30 °C for 2 h. The reaction was stopped by transferring the reaction mixtures to - 80 °C, followed by denaturation at 70 °C for 10 min. Further, after centrifugation, the supernatant was collected for subsequent LC-MS analysis, and 10 μL of the reaction was injected onto the chromatographic column. 3-methylglutaconyl-CoA compounds were analyzed using ion-pair reverse-phase HPLC. Mass spectrometric detection was carried out on an Agilent 6475 LC/TQ system (Agilent Technologies, Santa Clara, CA, USA) equipped with an AJS electrospray ionization (ESI) source operating in positive ionization mode.

### Data Analysis

Extracted-ion chromatogram peaks corresponding to 3-methylcrotonyl-CoA and 3-methylglutaconyl-CoA were integrated using MassHunter Qualitative Analysis software v10.0 (Agilent Technologies, Santa Clara, CA, USA), and the resulting peak areas were exported to spreadsheets and further processed in GraphPad Prism 10 (GraphPad Software, San Diego, CA).

### In-gel digestion and nano-LC-MS workflow

Proteins were separated by reducing SDS-PAGE and visualized with Coomassie Brilliant Blue. Gel bands were excised, diced to ∼1mm^3^, and subjected to in-gel tryptic digestion as described [16]. Briefly, gel pieces were washed with acetonitrile (ACN) and water, reduced with 2mM DTT, alkylated with 5mM iodoacetamide (30 min, RT, dark), and digested overnight at 37 °C with sequencing-grade trypsin. Peptides were extracted, dried, resuspended in 97:3:0.1 (H_2_O:ACN: Formic acid), and desalted using C18 StageTips. Peptide mixtures were analyzed by nanoLC-MS/MS on an UltiMate 3000 RSLCnano system coupled to a Q Exactive HF mass spectrometer (Thermo Fisher Scientific). Separation was achieved using a trap column (nanoEase M/Z Symmetry C18 (180 µm × 20 mm)) and an analytical column (nanoEase M/Z HSS C18 T3 (75 µm × 250 mm)) at 40 °C with a linear 85 min gradient from 1% to 40% mobile phase B (0.1% formic acid in acetonitrile) at 0.3 µl/min.

The mass spectrometer was operated in positive-ion mode with data-dependent acquisition. Full MS scans were recorded at a resolution of 60,000 at m/z 200, followed by up to 10 MS/MS scans per cycle at 15,000 resolution using higher-energy collisional dissociation with a normalized collision energy of 28. Dynamic exclusion was set to 20 s, and singly charged as well as unassigned ions were excluded from MS/MS analysis. Instrument performance was routinely assessed using standard quality control injections. Raw MS data were processed using MaxQuant (v1.6.17.0) with label-free quantification enabled and the match-between-runs function activated. Database searches were conducted against the UniProt reviewed protein database supplemented with common contaminants.

### Cryo-EM Grid Preparation and Data Acquisition

Cryo-EM grids were prepared by applying 3.5 µL of purified protein derived from membrane and cytosolic fractions onto glow-discharged gold Ultrafoil grids (R1.2/1.3, 300 mesh). Grids were blotted for 3.5 s using a blot force of 10 in a Vitrobot Mark IV (Thermo Fisher Scientific) operated at 4 °C and 100% relative humidity. Immediately after blotting, grids were plunge-frozen in liquid ethane for vitrification. Cryo-EM data from grids prepared with purified cytosolic fractions were collected on a Titan Krios G1 transmission electron microscope (Thermo Fisher Scientific) operated at 300 kV. The microscope was equipped with a Gatan K3 direct electron detector and a BioQuantum energy filter (Gatan) using a slit width of 20 eV. A total of 19,195 movies were recorded in electron counting mode using aberration-free image shift as implemented in EPU (Thermo Fisher Scientific). Data were acquired at a nominal magnification of 105,000 x, corresponding to a calibrated pixel size of 0.836 Å. Each movie consisted of 60 frames with a total electron dose of 60 e^−^/Å^2^ and a defocus range of −0.8 to −2.0 µm. Cryo-EM data from grids prepared with purified membrane fractions were acquired on a Glacios transmission electron microscope (Thermo Fisher Scientific) operated at 200 kV. The microscope was equipped with a Falcon 4i direct electron detector and a Selectris energy filter (Thermo Fisher Scientific) with a slit width of 20 eV. A total of 17,483 movies were collected in EER format at a nominal magnification of 130,000 x, corresponding to a calibrated pixel size of 0.69 Å. Each movie was recorded with a total electron dose of 60 e^−^/Å^2^ and a defocus range of −0.8 to −2.0 µm.

### Cryo-EM data processing

Cryo-EM data processing was performed using cryoSPARC v4.5.3 (Structura Biotechnology Inc.). Patch-based motion correction and contrast transfer function (CTF) estimation were applied to all micrographs [17]. Micrographs were manually inspected, and a subset was selected for initial particle picking. Manually picked particles were subjected to two-dimensional (2D) classification, and high-quality class averages were used as templates for automated particle picking. For the MCC complex purified from the membrane fraction (Supplementary Fig. 7), particles were extracted using a box size of 512 pixels and Fourier cropped to 168 pixels. Several rounds of 2D classification were performed to remove low-quality particles and artifacts. An initial ab initio reconstruction was generated from the selected particles, followed by heterogeneous refinement. Particles corresponding to the best-resolved classes were re-extracted with a box size of 512 pixels and Fourier cropped to 300 pixels, then subjected to non-uniform refinement [15] with D3 symmetry followed by local and global CTF correction, yielding a reconstruction at 2.9 Å resolution (Supplementary Fig. 2), as determined by gold-standard Fourier shell correlation using the 0.143 criterion. The final map was used to guide atomic model building.

For the MCC complex purified from the cytosolic fraction (Supplementary Fig. 8), particles were initially extracted using a box size of 480 pixels and Fourier cropped to 288 pixels. Multiple rounds of 2D classification were performed to eliminate junk particles. An initial ab initio reconstruction was generated, followed by heterogeneous refinement. Particles of overexpressed views were removed by using the ‘rebalance orientation’ job. Balanced particles were re-extracted with a box size of 480 pixels, Fourier cropped to 416 pixels and subjected to non-uniform refinement with D3 symmetry. This procedure produced a reconstruction with a final resolution of 2.2 Å (Supplementary Fig. 1), as determined by gold-standard Fourier shell correlation at the 0.143 cutoff. The resulting map was used for subsequent atomic model building.

### Model Building

To model the MCC complex (6AccA1:6AccD1) purified from both membrane and cytosolic fractions, an initial structural model was generated using CryFold prediction [18]. The predicted model was rigid-body fitted into the cryo-EM density map using ChimeraX [19], followed by iterative real-space refinement [20] in PHENIX [21]. Side chains were inspected visually and manually adjusted when supported by corresponding electron density. Flexible loops and terminal regions were built in Coot [22] wherever continuous and well-defined density was present. Residues lacking unambiguous side-chain density were modeled based on secondary structure geometry and stereochemical restraints. Regions without interpretable electron density were either assigned zero occupancy or omitted from the final model. All ligand molecules were placed into the cryo-EM density using Coot with monomer library restraints and subsequently refined in PHENIX. To model the covalent linkage between Lys564 of the biotin carboxyl carrier protein (BCCP) domain and biotin, a covalent bond restraint was defined in ChimeraX. A custom restraint describing the Lys-biotin covalent geometry was generated using the eLBOW module in PHENIX and applied during the final rounds of real-space refinement to maintain appropriate stereochemical parameters.

### Data and Material Availability

Cryo-EM maps and associated atomic models reported in this study have been deposited in the Electron Microscopy Data Bank under accession codes EMD-57132(MCC complex, BC-engaged BCCP state), EMD-57153 (MCC complex, CT-engaged BCCP state), and the Protein Data Bank with accession codes PDB_000029EX and PDB_000029GV, respectively.

## Acknowledgements

We thank Patrick Voskamp for technical laboratory support. We are grateful to Willem Noteborn and Ludovic Renault (Netherlands Centre for Electron Nanoscopy) for support with cryo-electron microscopy data collection. This work was supported by the Leiden Institute of Chemistry and by Oncode Accelerator, funded through the Dutch National Growth Fund (grant NGFPO2201). Cryo-electron microscopy data were collected at the Netherlands Centre for Electron Nanoscopy (NeCEN) as part of the Netherlands Electron Microscopy Infrastructure, funded through the National Roadmap for Large-Scale Research Infrastructure of the Dutch Research Council (project number 184.034.014).

## Author contributions

A.Y. and S.G. conceived the study. A.Y. and B.F. performed the experiments. A.Y. carried out biochemical analyses, sample preparation, grid screening, cryo-EM image processing, three-dimensional reconstruction, and model building. B.F. and A.Y. performed LC-MS experiments. Data analysis was performed by A.Y., B.F., and S.G. The manuscript was written by A.Y., B.F., and S.G., with input from all authors.

## Data availability

The data that support the findings of this study are available from the corresponding author upon request.

## References

1 Massey LK, Sokatch JR & Conrad RS (1976) Branched-chain amino acid catabolism in bacteria. Bacteriol Rev 40, 42–54.

2 Gallardo ME, Desviat LR, Rodríguez JM, Esparza-Gordillo J, Pérez-Cerdá C, Pérez B, Rodríguez-Pombo P, Criado O, Sanz R, Morton DH, Gibson KM, L. TP, Ribes A, de Córdoba SR, Ugarte M & Peñalva MÁ (2001) The Molecular Basis of 3-Methylcrotonylglycinuria, a Disorder of Leucine Catabolism. The American Journal of Human Genetics 68, 334–346.

3 Tong L (2013) Structure and function of biotin-dependent carboxylases. Cell Mol Life Sci 70, 863–891.

4 Chou C-Y, Yu LPC & Tong L (2009) Crystal Structure of Biotin Carboxylase in Complex with Substrates and Implications for Its Catalytic Mechanism *. Journal of Biological Chemistry 284, 11690–11697.

5 Mochalkin I, Miller JR, Evdokimov A, Lightle S, Yan C, Stover CK & Waldrop GL (2008) Structural evidence for substrate-induced synergism and half-sites reactivity in biotin carboxylase. Protein Science 17, 1706–1718.

6 Lietzan AD, Menefee AL, Zeczycki TN, Kumar S, Attwood PV, Wallace JC, Cleland WW & St. Maurice M (2011) Interaction between the Biotin Carboxyl Carrier Domain and the Biotin Carboxylase Domain in Pyruvate Carboxylase from Rhizobium etli. Biochemistry 50, 9708–9723.

7 Broussard TC, Pakhomova S, Neau DB, Bonnot R & Waldrop GL (2015) Structural Analysis of Substrate, Reaction Intermediate, and Product Binding in Haemophilus influenzae Biotin Carboxylase. Biochemistry 54, 3860–3870.

8 Ehebauer MT, Zimmermann M, Jakobi AJ, Noens EE, Laubitz D, Cichocki B, Marrakchi H, Lanéelle M-A, Daffé M, Sachse C, Dziembowski A, Sauer U & Wilmanns M (2015) Characterization of the Mycobacterial Acyl-CoA Carboxylase Holo Complexes Reveals Their Functional Expansion into Amino Acid Catabolism. PLOS Pathogens 11, e1004623.

9 Huang CS, Ge P, Zhou ZH & Tong L (2011) An unanticipated architecture of the 750 kD holoenzyme of 3-methylcrotonyl-CoA carboxylase. Nature 481, 219–223.

10 Hu JJ, Lee JKJ, Liu Y-T, Yu C, Huang L, Aphasizheva I, Aphasizhev R & Zhou ZH (2023) Discovery, structure, and function of filamentous 3-methylcrotonyl-CoA carboxylase. Structure 31, 100-110.e4.

11 Plaza-Pegueroles A, Aphasizheva I, Aphasizhev R, Fernández-Tornero C & Ruiz FM (2024) The cryo-EM structure of trypanosome 3-methylcrotonyl-CoA carboxylase provides mechanistic and dynamic insights into its enzymatic function. Structure 32, 930-940.e3.

12 Zhou F, Zhang Y, Zhu Y, Zhou Q, Shi Y & Hu Q (2024) Structural insights into human propionyl-CoA carboxylase (PCC) and 3-methylcrotonyl-CoA carboxylase (MCC). eLife 13.

13 Su J, Tian X, Cheng H, Liu D, Wang Z, Sun S, Wang H-W & Sui S-F (2025) Structural insight into synergistic activation of human 3-methylcrotonyl-CoA carboxylase. Nat Struct Mol Biol 32, 73–85.

14 Liang Y & Rubinstein JL (2025) Structural basis for substrate specificity and MSMEG_0435-0436 binding by the mycobacterial long-chain acyl-CoA carboxylase complex., 2025.10.28.685139.

15 Punjani A, Zhang H & Fleet DJ (2020) Non-uniform refinement: adaptive regularization improves single-particle cryo-EM reconstruction. Nat Methods 17, 1214–1221.

16 Shevchenko A, Tomas H, Havlis J, Olsen JV & Mann M (2006) In-gel digestion for mass spectrometric characterization of proteins and proteomes. Nat Protoc 1, 2856–2860.

17 Punjani A, Rubinstein JL, Fleet DJ & Brubaker MA (2017) cryoSPARC: algorithms for rapid unsupervised cryo-EM structure determination. Nat Methods 14, 290–296.

18 Su B, Huang K, Peng Z, Amunts A & Yang J (2024) Improved model building for cryo-EM maps using local attention and 3D rotary position embedding.

19 Pettersen EF, Goddard TD, Huang CC, Meng EC, Couch GS, Croll TI, Morris JH & Ferrin TE (2021) UCSF ChimeraX: Structure visualization for researchers, educators, and developers. Protein Sci 30, 70–82.

20 Afonine PV, Poon BK, Read RJ, Sobolev OV, Terwilliger TC, Urzhumtsev A & Adams PD (2018) Real-space refinement in PHENIX for cryo-EM and crystallography. Acta Crystallogr D Struct Biol 74, 531–544.

21 Liebschner D, Afonine PV, Baker ML, Bunkóczi G, Chen VB, Croll TI, Hintze B, Hung LW, Jain S, McCoy AJ, Moriarty NW, Oeffner RD, Poon BK, Prisant MG, Read RJ, Richardson JS, Richardson DC, Sammito MD, Sobolev OV, Stockwell DH, Terwilliger TC, Urzhumtsev AG, Videau LL, Williams CJ & Adams PD (2019) Macromolecular structure determination using X-rays, neutrons and electrons: recent developments in Phenix. Acta Crystallogr D Struct Biol 75, 861–877.

22 Emsley P, Lohkamp B, Scott WG & Cowtan K (2010) Features and development of Coot. Acta Crystallogr D Biol Crystallogr 66, 486–501.

