## Supplementary material for "Cryo-EM structures of mycobacterial MCC reveal carrier-domain translocation between catalytic sites": AccA1_AccD1_SI.pdf

**Supplementary Table 1.** List of Proteins Identified by LC–MS.

| Uniprot Id | No of peptide | MW (KDa) |
|---|---|---|
| I7G6G9_ AccA3 | 61 | 63 |
| I7G4Z0_ AccD5 | 45 | 58 |
| I7G857_ Pyruvate carboxylase | 25 | 120 |
| A0R1D8_AccA1 | 22 | 70 |
| I7GCS3_AccD1 | 20 | 54 |
| I7GGE0_AccD4 | 55 | 56 |
| I7F9I8_AccE5 | 8 | 10 |

**Supplementary Table 2.** Cryo-EM data collection, structure determination, model building, and refinement statistics.

| | BC engaged BCCP sate of MCC complex (EMD:57132,PDB: 29EX) | CT engaged BCCP sate of MCC complex (EMD:57153,PDB: 29GV) |
|---|---|---|
| **Data collection** | | |
| Microscope | Glacios | FEI Titan Krios |
| Voltage (kV) | 200 | 300 |
| Magnification | 130,000 | 105,000 |
| Electron exposure (e-/ Å$^2$) | 60 | 60 |
| Defocus range (μm) | 0.8-2.0 | 0.8-2.0 |
| Raw pixel size (Å) | 0.69 | 0.84 |
| Micrograph collected | 17,483 | 19,195 |
| **Reconstruction** | | |
| Symmetry | D3 | D3 |
| Initial particle images extracted (no.) | 280,017 | 50,17,262 |
| Initial particle images for 3D processing (no.) | 10,624 | 416,391 |
| Final Particle images(no.) | 6,673 | 177,311 |
| Map resolution (Å), FSC threshold (0.143) | 2.9 | 2.2 |
| FSC mask map resolution range (Å), FSC threshold (0.143) | 2.9-3.8 | 2.2-2.8 |
| Map sharpening B- factor | 45.2 | 60.0 |
| **Model Building and Refinement** | | |
| Initial model used | CryFold | CryFold |
| Model resolution (Å) | 3.1 | 3.6 |
| FSC threshold | 0.5 | 0.5 |
| CC (volume/mask) | 0.85/0.87 | 0.68/0.67 |
| Protein B factor (Å$^2$) | 51.21 | 82.24 |
| Ligand B factor (Å$^2$) | 20.00 | 20.00 |
| **Model composition** | | |
| Nonhydrogen atom | 46040 | 32186 |
| Protein residue | 6143 | 4289 |
| Ligands | BTN:6 | BTN:6 |
| **RMS deviations** | | |
| Bond length (Å) | 0.007 | 0.003 |
| Bond angle (°) | 0.597 | 0.523 |
| **Validation** | | |
| MolProbity score | 1.84 | 1.79 |
| Clashscore | 4.91 | 4.67 |
| Poor rotamers (%) | 3.22 | 3.54 |
| CaBLAM outlier(%) | 1.33 | 1.64 |
| **Ramachandran plot** | | |
| Favored (%) | 96.81 | 97.27 |
| Allowed (%) | 3.19 | 2.70 |
| Disallowed (%) | 0.00 | 0.02 |

**(a)** **(b)**

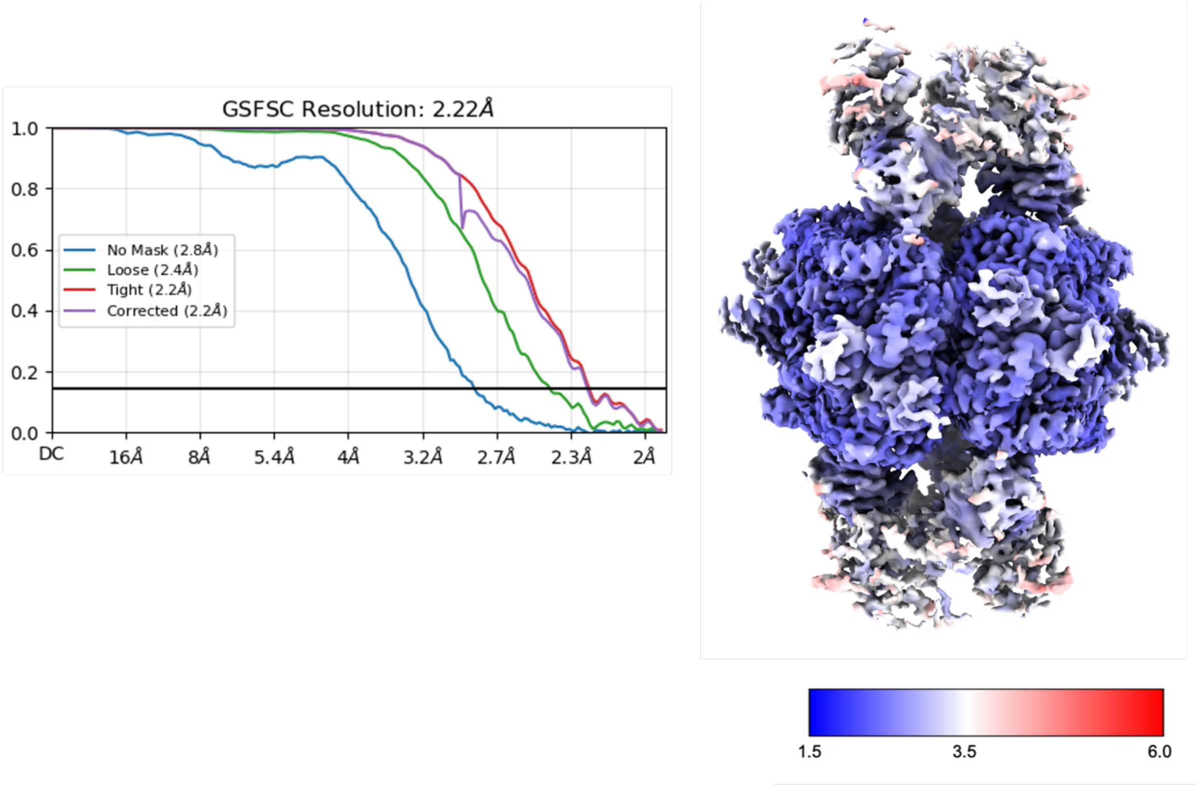


**Supplementary Fig. 1. a,** Fourier shell correlation (FSC) of the CT-engaged BCCP state of MCC complex. **b,** Estimated average resolution of the cryo-EM map.

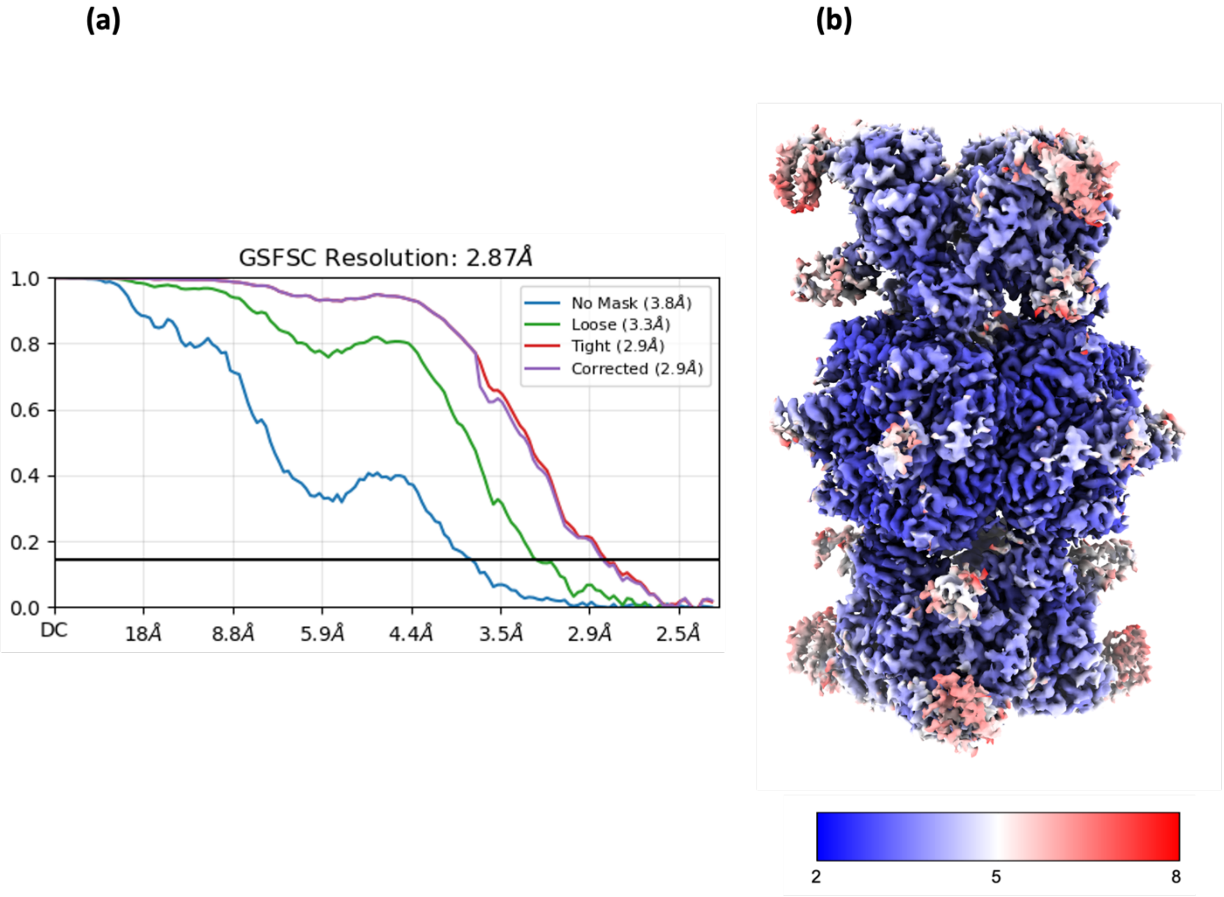


**Supplementary Fig. 2. a,** Fourier shell correlation (FSC) of the BC-engaged BCCP state of MCC complex. **b,** Estimated average resolution of the cryo-EM map.

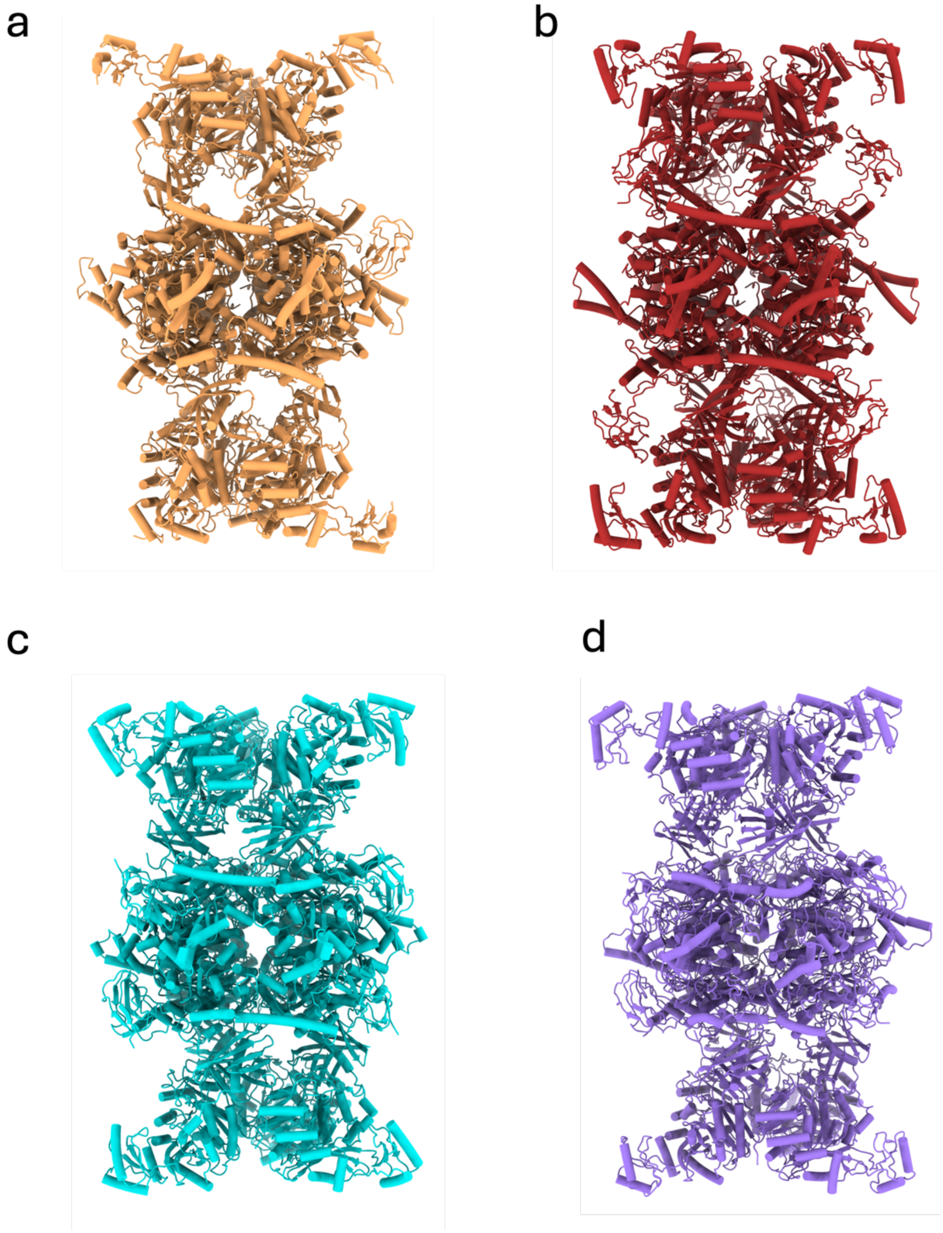


**Supplementary Fig. 3.** Reported cryo-EM structure of MCC complex from different species:
**a,** *P. aeruginosa* (PDB: 3U9S). **b,** *Homo sapiens* (PDB: 8JAK). **c,** *Trypanosoma brucei* (PDB: 8RTH). **d,** *Leishmania tarentolae* (PDB: 8F3D).

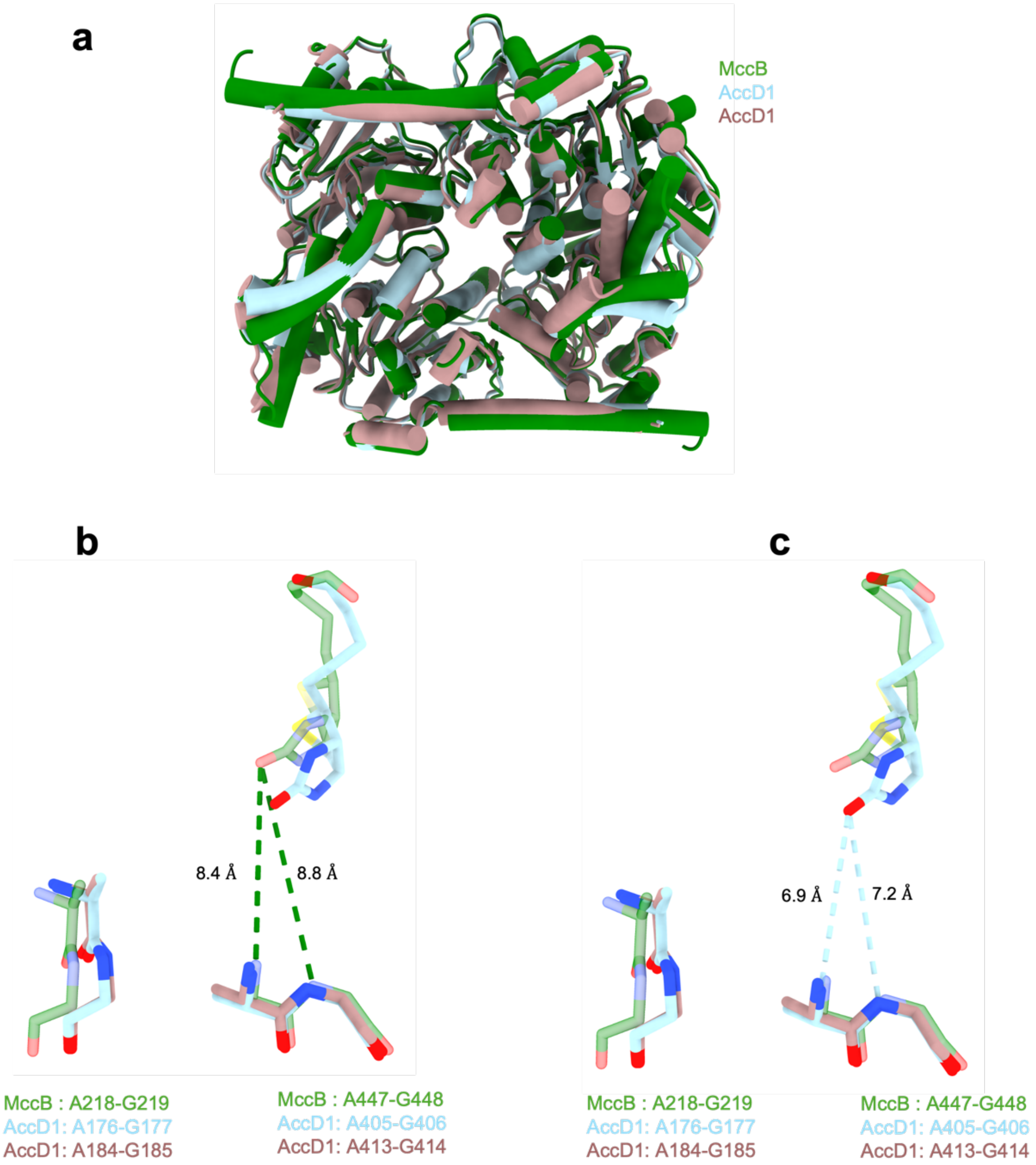


**Supplementary Fig. 4. Comparison of conserved oxyanion holes in the CT subunit of the MCC complex from human and *Mycobacterium smegmatis*, and from the hexameric AccD1 structure of *Mycobacterium tuberculosis*.**

**a,** Structural superposition of the MccB dimer from human (PDB: 8JAW, green) with the AccD1 dimer from *M. tuberculosis* (PDB: 4Q0G, brown) and the AccD1 dimer from *M. smegmatis* (light blue) in the CT-engaged BCCP state.

**b-c,** Enlarged views of the active sites after structural superposition, showing conserved oxyanion-hole residues (stick representation) in equivalent positions. **b,** Human MccB bound to biotin (green). **c** AccD1 from *M. smegmatis* bound to biotin (light blue). Distances between the biotin and the oxyanion-hole residues are indicated.

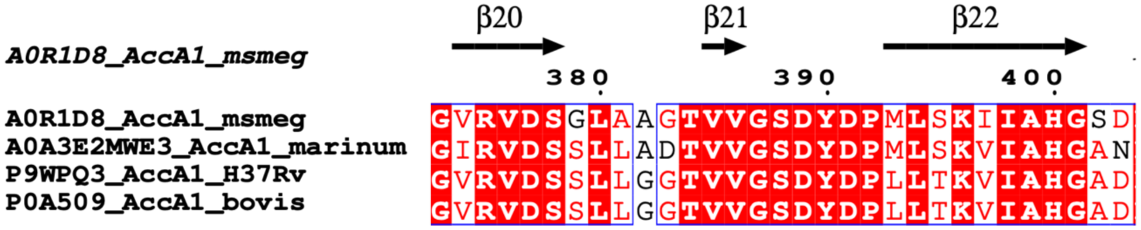


**Supplementary Fig. 5.** Sequence alignment of MCC α-subunit from different mycobacterium species (Only residues 373-403 are shown).

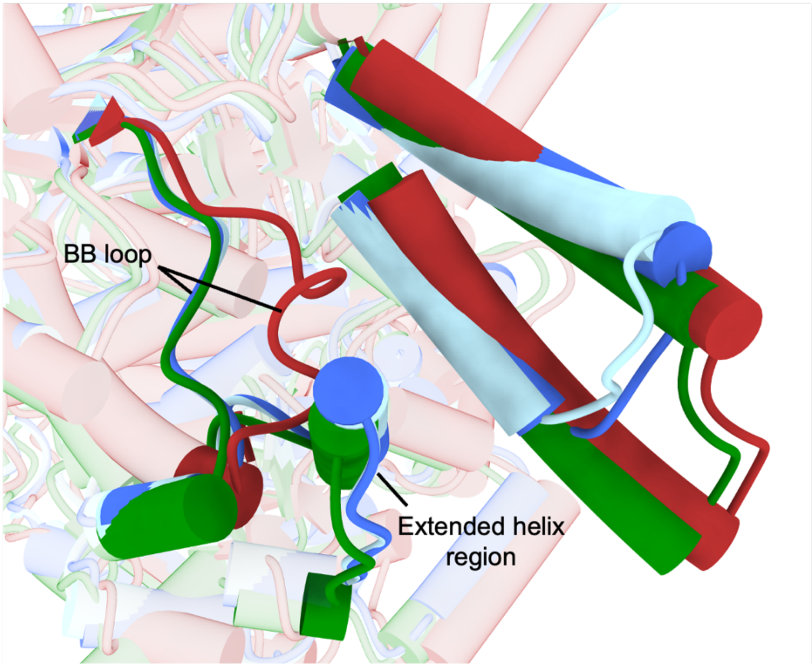


**Supplementary Fig. 6**. Structural superposition of the carboxyltransferase (CT) subunit in the BC- and CT-engaged BCCP states of the MCC complex (blue and light blue, respectively) with reported human MCC structures: the BC-engaged state (PDB: 8JAK, red) and the CT-engaged state (PDB: 8JAW, green). Conserved features near the BCCP-binding region are highlighted. The BB loop (blue and light blue) shows no significant conformational differences between the BC- and CT-engaged states.

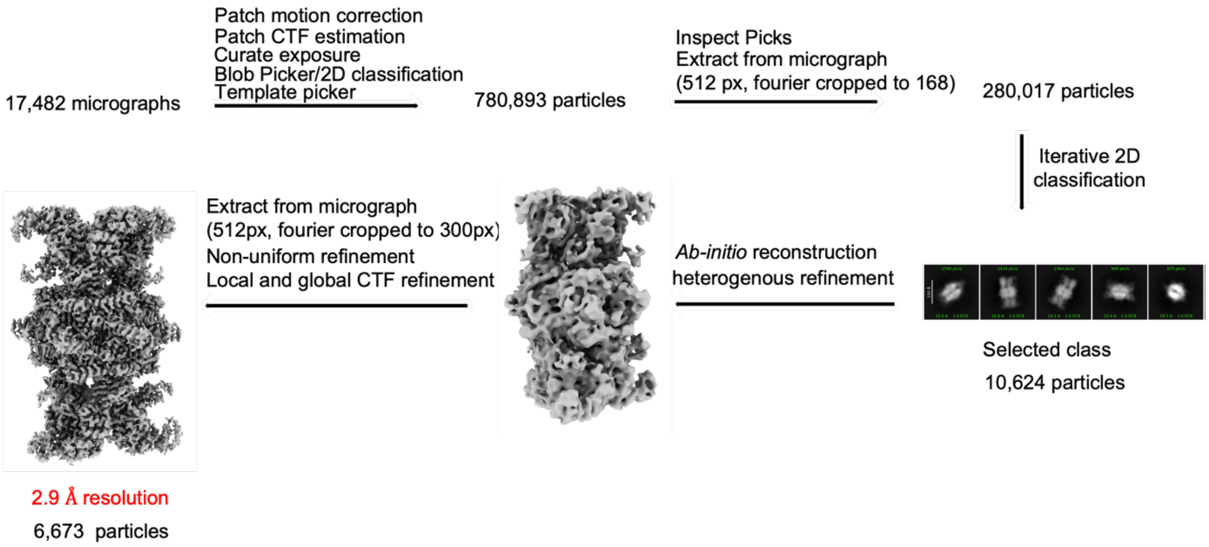


**Supplementary Fig. 7.** Image processing and classification workflow for the cryo-EM dataset for the BC-engaged BCCP state of MCC complex.

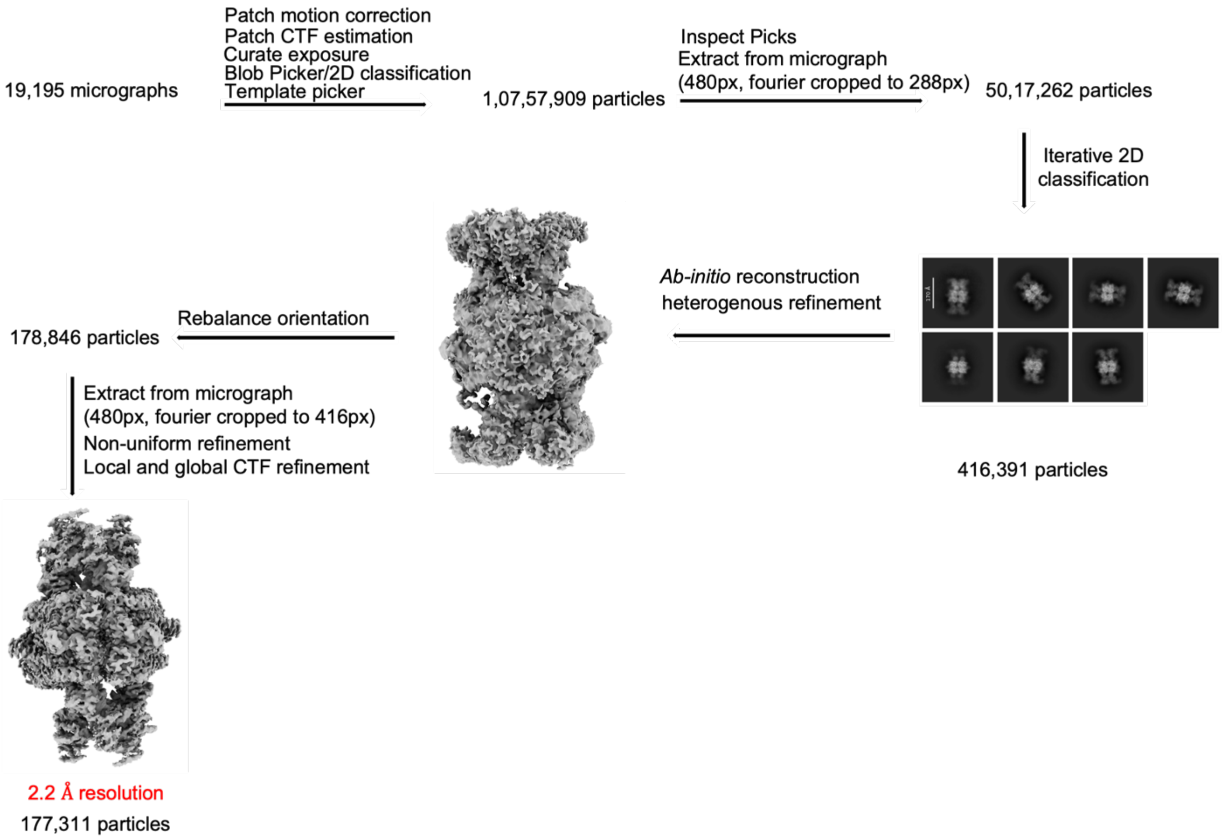


**Supplementary Fig. 8.** Image processing and classification workflow for the cryo-EM dataset for the CT-engaged BCCP state of MCC complex.
